# Exploring the mutational robustness of nucleic acids by searching genotype neighbourhoods in sequence space

**DOI:** 10.1101/091389

**Authors:** Qingtong Zhou, Xianbao Sun, Xiaole Xia, Zhou Fan, Zhaofeng Luo, Suwen Zhao, Haojun Liang, Eugene Shakhnovich

## Abstract

To assess the mutational robustness of nucleic acids, many genome- and protein-level studies have been performed; in these investigations, nucleic acids are treated as genetic information carriers and transferrers. However, the molecular mechanism through which mutations alter the structural, dynamic and functional properties of nucleic acids is poorly understood. Here, we performed SELEX in silico study to investigate the fitness distribution of the nucleic acid genotype neighborhood in a sequence space for L-Arm binding aptamer. Although most mutants of the L-Arm-binding aptamer failed to retain their ligand-binding ability, two novel functional genotype neighborhoods were isolated by SELEX in silico and experimentally verified to have similar binding affinity (K_d_ = 69.3 μM and 110.7 μM) as the wild-type aptamer (K_d_ = 114.4 μM). Based on data from the current study and previous research, mutational robustness is strongly influenced by the local base environment and ligand-binding mode, whereas bases distant from the binding pocket provide potential evolutionary pathways to approach global fitness maximum. Our work provides an example of successful application of SELEX in silico to optimize an aptamer and demonstrates the strong sensitivity of mutational robustness to the site of genetic variation.

---

As one of the most important biological macromolecules, nucleic acids have diverse functions in encoding, transmission and expression of genetic information. This diversity is due to the vast sequence space of nucleic acids. High dimensionality of sequence space provides multiple evolutionary pathways to evolve specific phenotypes under selection pressure. However such pathways might pass through local fitness minima (valleys on fitness landscape) due to detrimental effects of mutation in immediate vicinity of evolved genotypes. To address the fitness effect of mutations, extensive studies have focused on understanding mutational robustness at the genome and protein levels. Previous analyses of DNA sequencing data and mutation accumulation and mutagenesis experiments have revealed that more than 90% of gene knockouts in Escherichia coli are nonlethal,^1^ whereas in humans, most amino acid substitutions^2^ have fitness effects, amounting to selection coefficients, in the range of 10^−3^ to 10^−1^, and relatively few substitutions have effects greater than 0.1. The significant mutational robustness of cellular organisms could be explained by buffering mechanisms, including alternative metabolic pathways, genetic redundancy, and modularity. In a biological system without a buffering mechanism, such as some RNA viruses,^3,4^ random nucleotide mutations can reduce fitness by an average of nearly 50%, with up to 40% mutations being lethal. These numbers are similar to those found for DNA viruses,^5^ and both of these viruses exhibit greater mutational sensitivity than cellular organisms. The role of nucleic acids is genetic information carriers and transferrers, but the in-depth mutational robustness of nucleic acids themselves, i.e., how mutations alter the structural, dynamic and functional properties of nucleic acids, remains poorly understood. The exploration of nucleic acid sequence space is largely limited by the available experimental technologies. However their reach is not sufficient to cover the vast sequence space limiting the extent to which the mutational robustness of functional nucleic acids can be explored. Therefore, a comprehensive molecular-level analysis of the mutational effects on the structural, dynamic and functional properties of nucleic acids will provide a solid basis for understanding of molecular evolution of nucleic acids.

A nucleic acid can adopt distinct secondary or folded tertiary structures that bind targets potently and selectively. These structures, which are denoted aptamers, are generally identified from a random sequence library using Systematic Evolution of Ligands by Exponential Enrichment (SELEX).^6–8^ Through the efforts of many researchers, SELEX technology has evolved rapidly, and the current technology enables the identification of a wide range of aptamer targets, ranging from small molecules and metal ions to proteins, biological cells, and tissues. One variant of SELEX is genomic SELEX,^9^ which aims to identify genome-encoded nucleic acids with defined properties from a library consisting of short fragments from the human genome rather than random sequences. Through genomic SELEX, both ATP-binding aptamers and GTP-binding motifs were found to be encoded in genomic sequences,^10,11^ which provides an interesting perspective on gene regulation. Taking the target-binding affinity as the fitness indicator, aptamers are fitness peaks in a sequence space and are surrounded by many genotypes some of which have the ability to bind targets.^12,13^ A 40-mer aptamer has 120 single-mutant, 7020 double-mutant, and 266,760 triple-mutant genotype neighborhoods in the sequence space (consisting of 4^40^ ≈ 10^24^ sequences). Although the initial SELEX library comprises up to 10^18^ sequences, it remains very difficult for SELEX to effectively identify functional sequences from an aptamer genotype neighborhood (Supplementary Figure S1). Limited by a lack of high-throughput and parallel experimental technologies,^14–18^ an exhaustive search is extremely difficult. Thus, although the problem of finding a functional aptamer in a sequence space has been successfully addressed by SELEX, the greater challenge is to optimize the found aptamers towards better ligand-binding affinity or selectivity, i.e., the inference of local fitness maximum to global fitness maximum in a sequence space, seems unsolvable. Consequently, the mutational robustness of nucleic acids is not fully understood.

Computational approaches are rapid, efficient and parallelizable and have thus become important tools in nucleic acid research.^21–27^ In our previous work,^28^ we proposed a computational approach involving the application of SELEX in silico for aptamer selection and successfully identified six novel theophylline-binding RNA aptamers from 4^13^ sequences. In the present study, we selected the L-argininamide (L-Arm)-binding aptamer (the first solved 3D structure of a DNA aptamer^29^) as our research system and used SELEX in silico to predict the fitness (defined as the ligand-binding affinity K_d_) of each aptamer genotype neighborhood in the sequence space (Supplementary Text S1). The L-Arm-binding aptamer consists of a stem region (bases ^1–7^ and 18–24) and a non-canonical region (bases 8–17), which form the binding pocket^19,29–31^(Figure 1B). Base C9, which is stacked by a reversed Hoogsteen mismatch pair, A8–C17, and a Watson– Crick pair, G10–C16, forms two hydrogen bonds with L-Arm on its Watson–Crick edge. In the current study, we focused on the mutations in the binding pocket with the exception of C9, i.e., on the mutations in bases 8 and 10–17. All of the mutants (4^9^ = 262144) were analyzed at the stage of secondary structure analysis, whereas at the molecular dynamic (MD)-based virtual screening stage, the mutants with Hamming distances ranging from 1 to 3 to the original aptamer (2619 mutants in total) were selected for in depth analysis. Two novel functional genotype neighborhoods of L-Arm-binding aptamers were identified through SELEX in silico to exhibit comparable fitness (experimental K_d_ = 69.3 μM and 110.7 μM)) to the wildtype (WT) aptamer (experimental K_d_ = 114.4 μM). Combined with previously reported data,^19^ the constructed fitness landscape suggests that the mutational robustness of nucleic acids is generally low but infrequently high in certain evolutional direction. The target-binding ability of nucleic acids is extremely sensitive to the sequence variation in or near the binding pocket as expected, whereas bases distant from the binding pocket exhibit considerable tolerance to substitutions and represent a potential evolutional pathway for approaching the global fitness maximum.

**Table 1:**
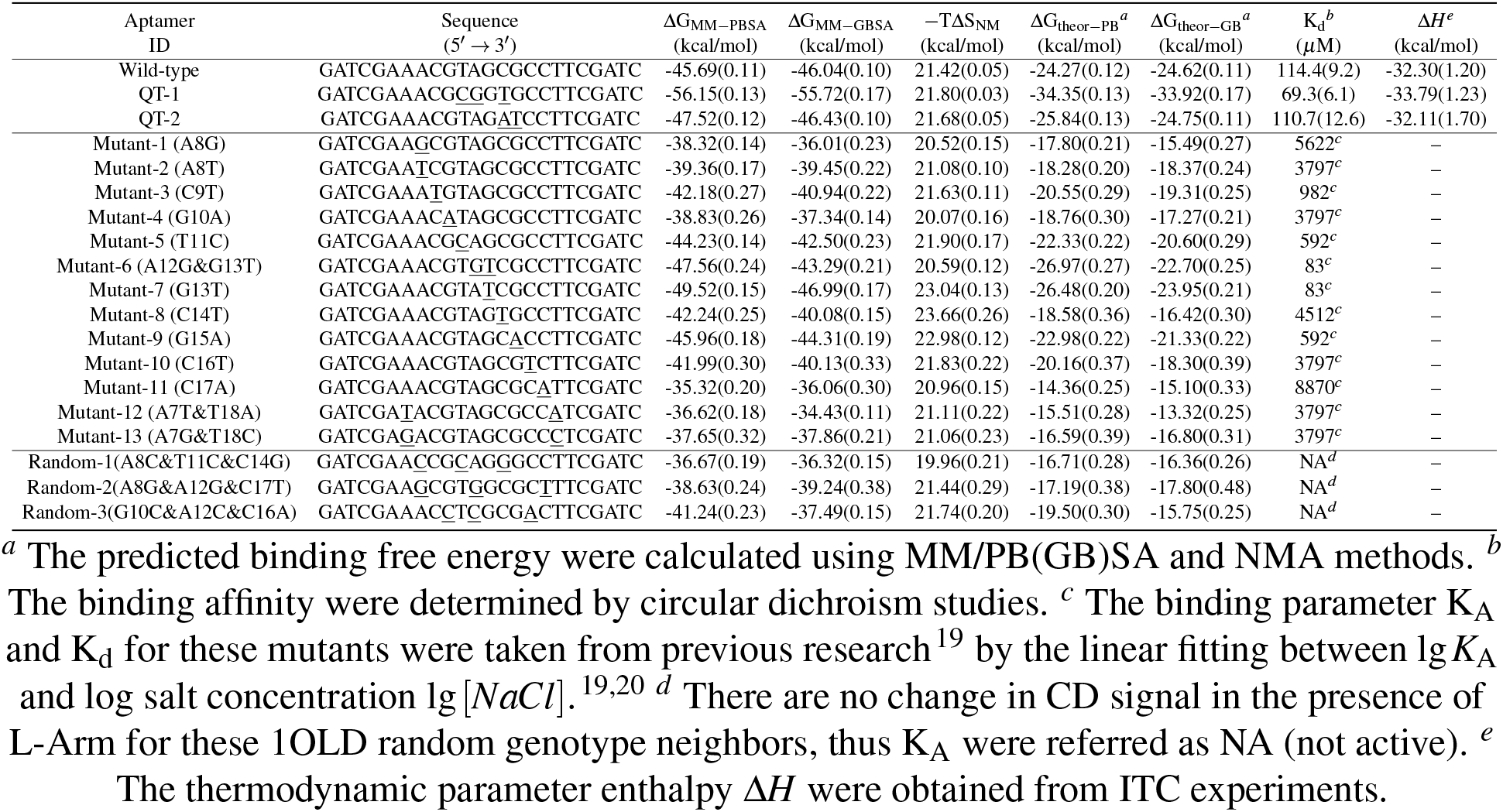
Binding affinity estimates for the wildtype L-Arm-binding aptamers and its genotype neighborhoods.

**Figure 1:**
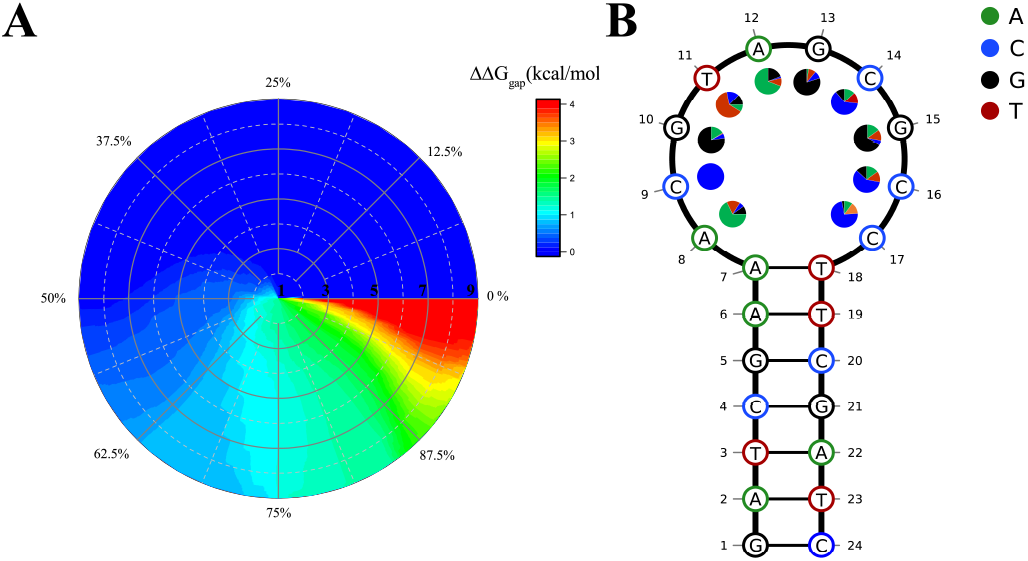
(A) Distribution of the free energy gaps in sequence space. The center of the polar plot is the WT L-Arm-binding DNA aptamer, the distance from the center indicates the corresponding Hamming distance of the mutants, the angle indicates the proportion of target motif foldable sequences in each sequence subspace, and the color represents the corresponding free energy gap (ΔG_gap_). (B) Secondary structure of the WT aptamer. The base preferences at each position in the non-canonical region were calculated for the screened best 100 sequences, which were selected by SELEX in silico from the 2619 closest neighbours (whose Hamming distance to the WT aptamer is no greater than three).

The minimum free energy (MFE) secondary structures of the 262144 mutants can be grouped into 57 unique structural motifs, among which the L-Arm-binding motif (the target motif identified by SELEX in silico, Figure 1B) is the most populated (118,127 sequences). All of the mutants can fold into the target motif with varying energy penalties (Supplementary Figure S2), and an average free-energy gap ΔΔG_gap_ (the difference between the lowest secondary structure energy state ΔG_MFE_ and the target secondary structure state ΔG_target,_ defined as ΔG_MFE_−ΔG_target_) of 0.78 kcal/mol was obtained. Surprisingly, 84% of the mutants (220,721 sequences) have a lower ΔΔG_gap_ than that of the WT aptamer (1.44 kcal/mol). As shown in Figure 1A, the distribution of the ΔΔG_gap_ in the sequence subspace (each sequence subspace was composed of mutated sequences with the same Hamming distance to the original aptamer thus contained 3^n^C^n^_9_ sequences, where *n* is the Hamming distance) was found to be consistent, regardless of the value of n, indicating that the secondary structure of the DNA aptamer exhibits remarkable tolerance to base substitutions. This finding is different from that for the theophylline-binding RNA aptamer,^28^ which has a complex secondary structure and is very sensitive to base substitutions, presented as a sharp peak on a rugged landscape. Similar to SELEX,^28^ MD-based virtual screening approaches bias the initial library toward ligand binding by predicting the ligand-binding free energy, and after several rounds of sequence enrichment, the potential aptamers are enriched. After two rounds of MD-based virtual screening, 100 of the 2370 sequences remained due to their high stability or low binding free energy, and the base preferences at each position were then calculated (Figure 1B). The percentage of the most populated bases ranged from the highest peak at the 13^th^ base (cytosine, 80%) to the lowest peak at the 16^th^ base (cytosine, 59%), whereas the reference values for the original and substituted bases among the 2619 mutants were approximately 68.3% and 10.6%, respectively. Although only the closest genotype neighborhood in the sequence space (Hamming distance less than 3) was searched in the current study, the mutational effect appears highly position-dependent. At positions 10, 13, and 17, the original base became more dominant, whereas multiple mutations of the 14^th^ or 16^th^ base were allowable.

As shown in Table 1 and Figure 2, ensembles of 20 simulations were run to obtain sufficient sampling of the conformational space,^32^ and the collected L-Arm-DNA complex snapshots were then subjected to MM/PB(GB)SA calculations and normal mode analysis (NMA) to estimate the enthalpic and entropic contributions to the binding free energy, respectively. The snapshot-based normalized frequency distributions of ΔG_MM=PBSA_, ΔG_MM=GBSA_ and − TΔS^NM^ presented welldefined Gaussian distributions (Supplementary Figure S3-S5). The calculated binding free energy of the WT aptamer genotype neighbours that reported in previous research^19^ agreed with their experimental mutational effect. Compared with the WT aptamer, most mutants have significantly higher calculated binding free energy, which are correctly predicted to bind ligand with lower binding affinity as revealed by experiment.^19^ Interestingly, two mutants Mutant-6 (A12G&G13T) and Mutant-7 (G13T) with slightly shifted to the left normalized frequency distribution of ΔG_MM-PBSA_ toward lower binding free energy (Figure 2A), could bind ligand tighter than the WT aptamer. Surprisingly, in silico selected genotype neighborhood aptamer QT-1, the best one predicted by SELEX in silico, has lower predicted binding free energy (the mean ΔG_MM=PBSA_ was −56.15 kcal/ mol) than the WT aptamer (−45.59 kcal/mol), while that of aptamer QT-2 was −47.52 kcal/mol. Similar effects were observed for ΔG_MM=GBSA_: aptamer QT-1 was the strongest, followed by aptamer QT-2 and the WT aptamer. The calculated entropies, −TΔS_NM_, of these three aptamers have coinciding mean (approximately 21.6 kcal/mol) and standard deviation values. Thus, two novel sequences (QT-1 and QT-2), which were identified through SELEX in silico from the aptamer closest neighborhood, were predicted to bind L-Arm as potently as the WT aptamer, and this finding was further experimentally verified (Supplementary Text S2).

**Figure 2:**
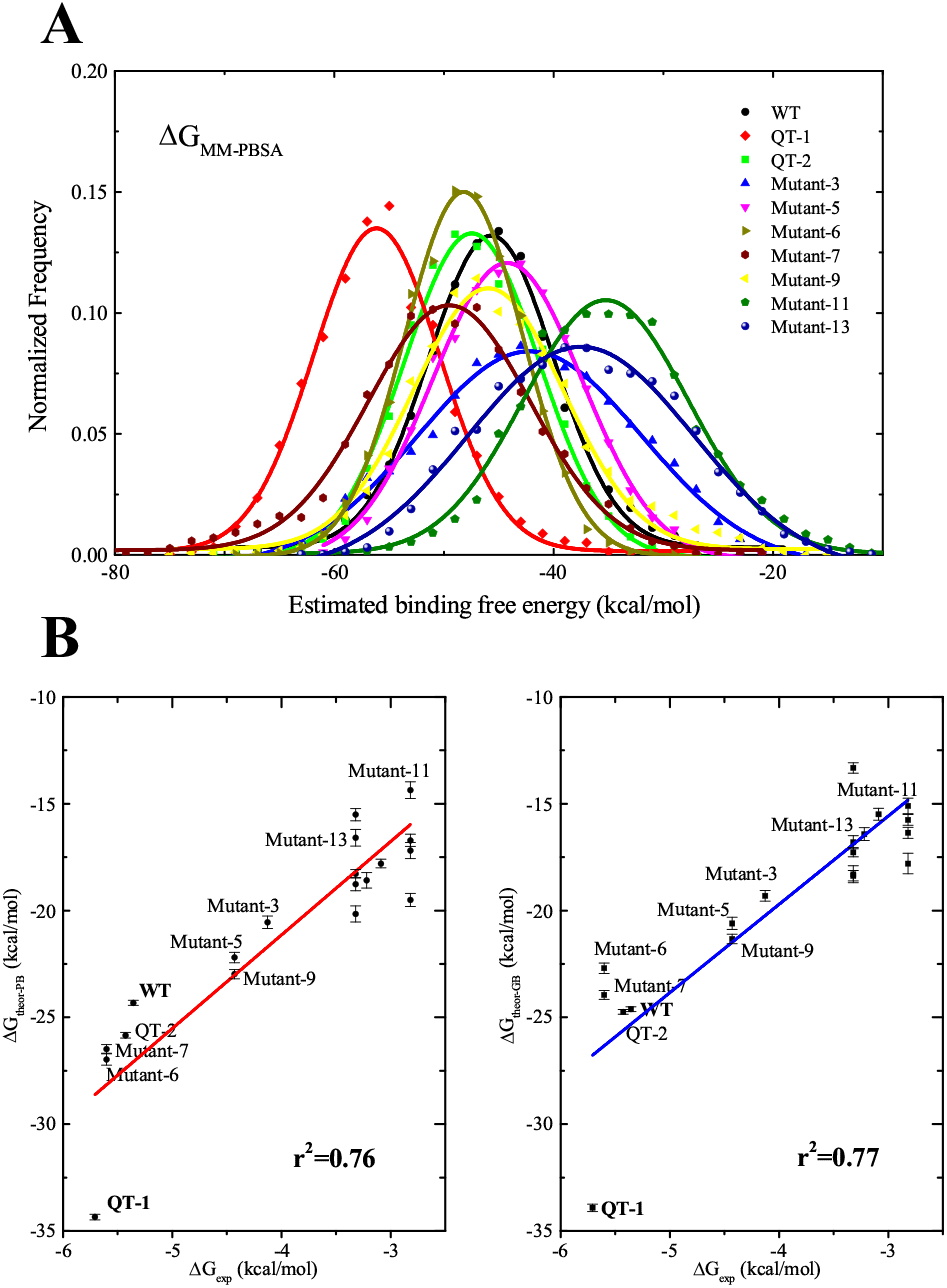
(A) Normalized frequency distribution of the calculated binding free energies. Distributions for MM/PBSA (ΔG_MM-PBSA_) are shown in per snapshot for the WT L-Arm-binding aptamers and its genotype neighborhoods. The expected normal distribution given the same mean and standard deviation for each data set is shown by the lines. (B) Comparison between the experimental ΔG_exp_ (kcal/mol)) and the calculated ΔG_theor-PB_ (left) and ΔG_theor-GB_ (right) using MM/PB(GB)SA and normal-mode analysis. Error bars show the standard errors. The line represents a linear regression performed on each data set.

**Figure 3:**
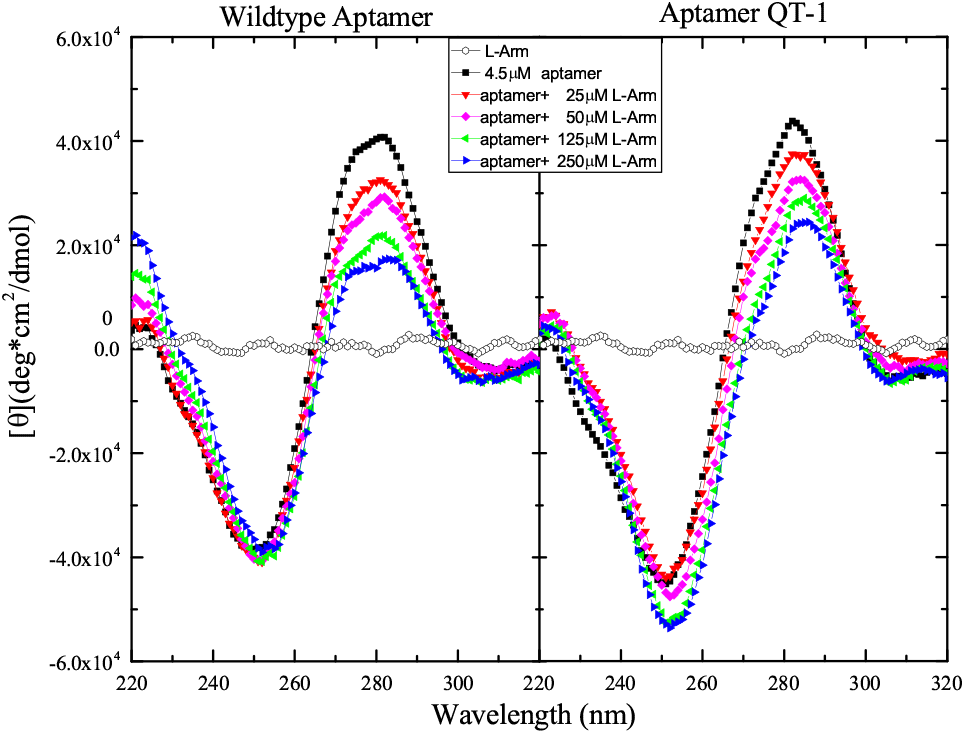
Circular dichroism (CD) spectra of the 4.5 μM L-Arm-binding DNA aptamers titrated with various concentrations of L-Arm in 10 mM sodium phosphate, 25 mM NaCl, pH 6.5. (Left) the WT aptamer; (Right) In silico screened aptamer QT-1;

Circular dichroism (CD) has been extensively used in research on nucleic acids because of its sensitivity to the conformation of anisotropic molecules.^19,33–35^ As shown in Figure 3, CD spectra were recorded by titrating the DNA aptamer at various concentrations of L-Arm. The WT aptamer displayed a positive peak at 280 nm in the CD spectra, whereas increasing concentrations of L-Arm decreased the molar ellipticity in this region (270–290 nm). This intensity change could indicate that the aptamer has changed its conformation to bind to the ligand, known as the induced-fit binding mechanism.^30,36^ The QT-1 and QT-2 sequences exhibit similar changes in the CD spectra, which demonstrates that these sequences may bind L-Arm in a manner similar to that found for the WT aptamer. Conversely, for many randomly selected genotype neighborhoods and previously reported clone 12-28 mutants,^19^ no changes in the CD signal were found in the presence of L-Arm, which is consistent with their extremely low ligand-binding affinity. To calculate the dissociation constant of the binding (K_d_), we analyzed the CD spectra using the optical curve direct fitting method. 24, 28, 29 (Supplementary Figure S6) For the WT aptamer, the value of K_d_ was 114.4±9:2 μM, which is similar to previously reported (~100μM^19^ and 134.6μM^33^for the longer 28-mer aptamer, 165.7μM^34^for 24-mer 1OLD aptamer). The K_d_ of the QT-2 aptamer is similar to that of the WT aptamer (110.7±12:6 μM), whereas the QT-1 aptamer exhibited strongest binding affinity with L-Arm (69.3±6:1 μM), which is generally consistent with our computational prediction. To obtain the enthalpic contribution to the ligand-binding process, we performed “model-free” isothermal titration calorimetry (ITC) studies^34,37^ to avoid any possible fitting bias (Supplementary Figure S7). By integrating the corrected area under the peaks, the overall enthalpy of binding for the WT enthalpy was found to equal −32.30±1.2 kcal/mol. In contrast, the QT-1 aptamer has a lower ΔH (−33.79±1.23 kcal/mol) than that of QT-2 (-32.11±1.7 kcal/mol). Comparing the experimental data and our prediction (Figure 2B), the coefficients of determination *r^2^* (0.76 for ΔG_theor−PB_ and 0.77 for ΔG_theor−GB_)were obtained, suggesting that MM/PB(GB)SA and NMA methods can accurately rank the ordering of ligand binding affinity of the mutants around WT aptamer. As noted, the overall entropy change in a binding system^33,38–40^ is a combination of aptamer conformational changes, re-organization of the solvent environment, changes in the translational and conformational freedom of the ligand, and the release of counterion molecules. The development of binding free energy calculation technologies especially entropy estimation methods will greatly facilitate fast and accurate selection of functional nucleic acid sequence from the vast sequence space.

The QT-1 aptamer is a triple mutant (T11C&A12G&C14T) of the WT aptamer, whereas QT-2 is a double mutant (C14A&G15T. As observed from the binding conformations, these mutations did not change the overall structure of the aptamer binding pockets. As shown in Figure 4, the guanidinium end of L-Arm was directed toward C16-C17 and forms two hydrogen bonds with the Watson-Crick edge of C9 of both the WT aptamer and its genotype neighborhood QT-1. The guanidinium-C9 pair was further stacked by a Watson-Crick G10•C16 pair. For the WT aptamer, the peptide linkage of L-Arm was directed toward A12 and forms a hydrogen bond with the sugar phosphate backbone of G10 and G13. However, the T11C mutation in the QT-1 aptamer weakens the occasional contact within T11-G15 and facilitates the folding of the T11-G15 loop segment toward L-Arm. The A12G mutation in particular successfully introduces an additional interaction between L-Arm and the carbonyl at C6 of guanine G12. These favorable interactions induced by mutations are conducive to the binding of L-Arm with the QT-1 aptamer. Similarly, the two mutations in aptamer QT-2 are relatively far from the binding pockets and appear to act as neutral mutations.

**Figure 4:**
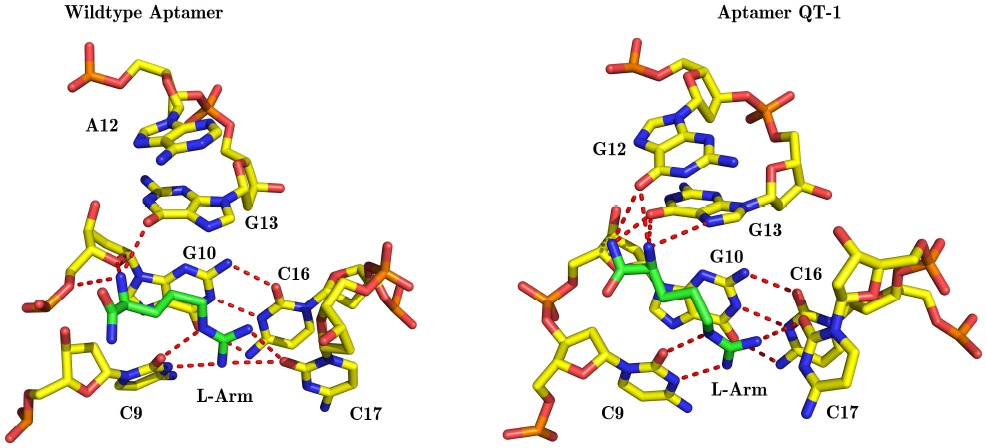
Comparison of the binding modes of L-Arm with the WT aptamer (Left) and in silico screened aptamer QT-1 (Right). The color of carbon in the aptamer was set to yellow, while for L-Arm it was blue. The red dashed lines indicate hydrogen-bonding interactions. Different from the WT aptamer by only three bases, the in silico screened aptamer QT-1 stabilizes L-Arm by constructing a closer binding pocket and forming extra hydrogen bonds between the base edges of G12 and L-Arm.

Based on earlier data^19^ and the current study (Supplementary Table S1), the fitness landscape was constructed to reflect how mutations alter the nucleic acid ligand-recognition ability.^41,42^ As shown in Figure 5, the WT aptamer is located at the origin of the x-y plane, whereas the mutations that occurred in the non-canonical region were represented at different coordinate azimuths to the positions of the mutated bases. The height of each mutant was represented by its ligand binding constant K_A_ with a corresponding color. Surrounding the WT aptamer, there are one single mutant G13T, two double mutants (A12G&G13T and C14A&G15T) and a triple mutant (T11C&A12G&C14T) with equivalent fitness. Unsurprisingly, almost all of the mutations around the binding pocket (A8T, A8G, C9T, G10A, C16T, C17A, A7T&T18A, and A7G&T18C) abolished the binding, which suggested that these bases were conserved for L-Arm binding and display significant low mutational tolerance. This conclusion is reasonable because each base surrounding the pocket plays an indispensable role in maintaining the particular ligand binding mode as follows: C9 is the partner of the hydrogens bonds for the ligand, the Watson-Crick pair G10•C16 and reverse A8•C17 can stack, and the Watson-Crick pair A7•T18 is the connector of the stem region and the non-canonical region of the aptamer. However, the bases located far from the binding pocket (T11, A12, G13, C14, and G15) show remarkable tolerance to mutations. The stepwise mutations G13T, A12G, and T11C&T13G&C14T can successfully evolve the WT aptamer to QT-1 without any fitness loss, which establishes an evolutional neutral pathway from one fitness peak to another higher fitness peak through local exploration. Thus, from a macroscopic perspective, functional aptamers are rare and evolutionarily isolated from one another in the sequence space, and the fitness landscape is a rugged “Badlands” landscape with multiple peaks.^12,13,19,28^ Benefit from the huge screened nucleic acid sequence library (up to 10^18^ sequences) and enrichment of ligand-binding nucleic acid, SELEX technology has greatly increased the observed probabilities of fitness peaks in sequence space. From a microscopic point of view, the majority of the mutants will lose their fitness, whereas only a few genotype neighborhoods in certain regions could be functional. The resulting fitness landscape is Fujiyama-like, and SELEX in silico could be adopted for its detailed exploration.

**Figure 5:**
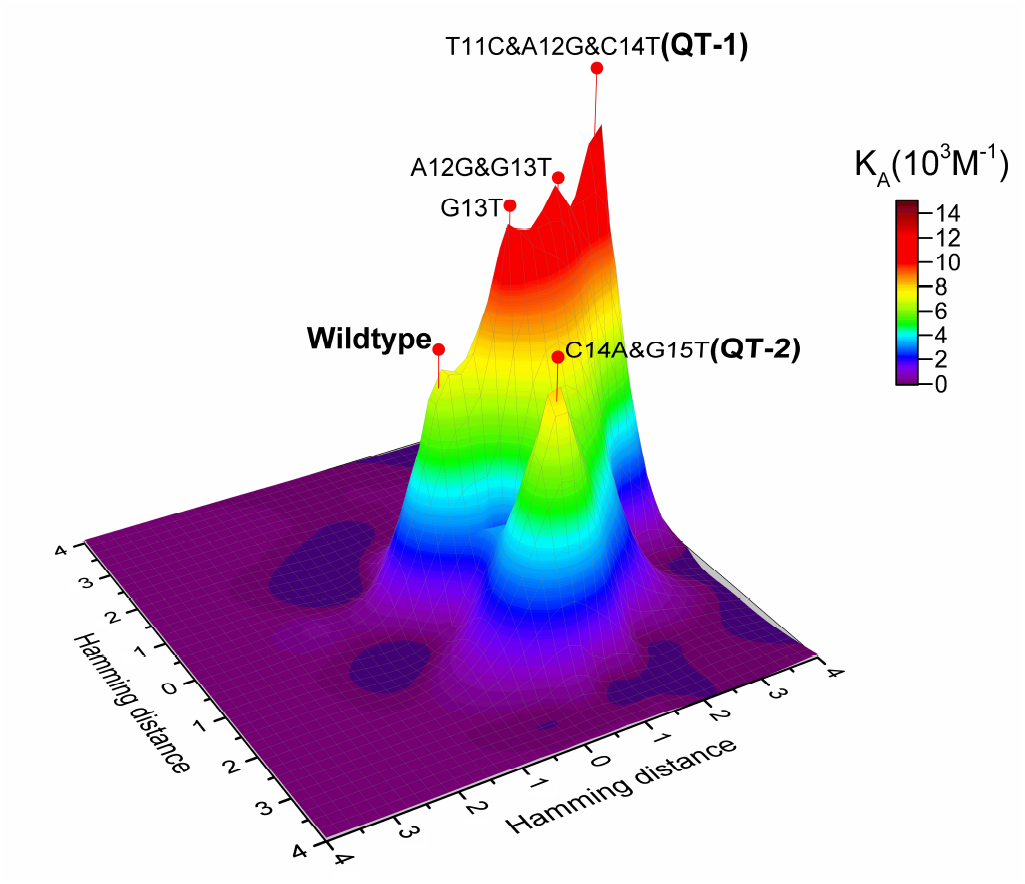
Experimental nucleic acid fitness landscape of ligand binding. Thirteen mutants from a previous work^19^ and ten mutants in this work were used to construct the fitness landscape. The intensity of the fitness peak was represented by the binding constant K_A_ with corresponding height and color.

In summary, we applied SELEX in silico to investigate the fitness distribution of nucleic acid genotype neighborhoods in a sequence space. Most mutants fail to bind the ligand with sufficient affinity, which is consistent with previous research for L-Arm binding DNA aptamer^19^ and other aptamers,^12,14,22,28^ indicates that the aptamer is very resistant to base substitutions and relies on the local sequence environment for target binding. Two novel aptamers were experimentally verified to exhibit similar fitness as the WT aptamer. The experimental nucleic acid fitness landscape constructed based on the current work and previous research^19^ suggests that the mutational robustness of nucleic acids is generally low but infrequently high in certain evolutional direction. Our work provides an example of successful application of SELEX in silico for aptamer optimization and demonstrates the complexity of the mutational robustness of nucleic acids from a novel perspective.

## Acknowledgement

The computations in this paper were run on the Odyssey cluster supported by the FAS Division of Science, Research Computing Group at Harvard University, the Shanghai Supercomputer Center and the Supercomputing Center of USTC. This work was supported by the China Postdoctoral Science Foundation [2016M591721 to Q.Z.], the National Natural Science Foundation of China [51573175, 91427304, 21434007 to H.L.] and the National Institutes of Health [GM068670 to E.S.].

## Supporting Information Available

Computational Details, Experimental Procedures, Supporting Figures and Tables. This material is available free of charge via the Internet at http://pubs.acs.org/.

## Graphical TOC Entry

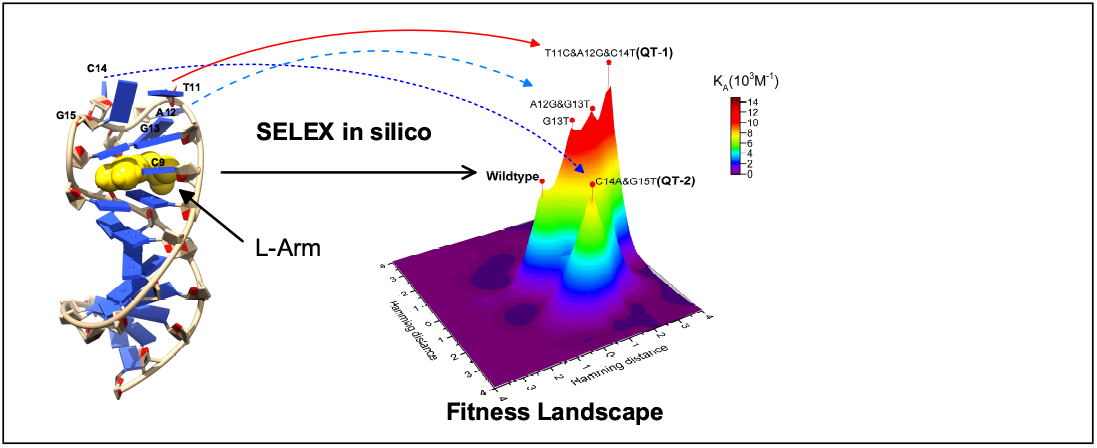

